# Genomic prediction accuracies and abilities for growth and wood quality traits of Scots pine, using genotyping-by-sequencing (GBS) data

**DOI:** 10.1101/607648

**Authors:** Ainhoa Calleja-Rodriguez, Jin Pan, Tomas Funda, Zhi-Qiang Chen, John Baison, Fikret Isik, Sara Abrahamsson, Harry X. Wu

**Affiliations:** Skogforsk (the Forestry Research Institute of Sweden), Box 3, SE-918 21 Sävar, Sweden; Umeå Plant Science Centre, Department of Forest Genetics and Plant Physiology, Swedish University of Agricultural Science, SE-901 83 Umeå, Sweden; Department of Genetics and Breeding, Faculty of Agrobiology and Natural Resources, Czech University of Life Sciences Prague, Kamýcká 129, 165 00 Prague, Czech Republic; Key Laboratory of Forest Genetics and Biotechnology, Nanjing Forestry University, Nanjing, 210037 China; Department of Forestry and Environmental Resources, North Carolina State University, Raleigh, NC 27695, USA; BAICTBMD, Beijing Forestry University, Beijing, 100083 China; NRCA, CSIRO, Canberra, ACT 2601, Australia

**Author notes:** Correspondence to: Harry X. Wu, Umeå Plant Science Centre, Department of Forest Genetics and Plant Physiology, Swedish University of Agricultural Science, SE-901 83 Umeå, Sweden.

**Keywords:** Scots pine, genotyping-by-sequencing, genomic relationship, GBLUP, Bayesian ridge regression, Bayesian-LASSO

## Abstract

Higher genetic gains can be achieved through genomic selection (GS) by shortening time of progeny testing in tree breeding programs. Genotyping-by-sequencing (GBS), combined with two imputation methods, allowed us to perform the current genomic prediction study in Scots pine (*Pinus sylvestris* L.). 694 individuals representing 183 full-sib families were genotyped and phenotyped for growth and wood quality traits. 8719 SNPs were used to compare different genomic prediction models. In addition, the impact on the predictive ability (PA) and prediction accuracy to estimate genomic breeding values was evaluated by assigning different ratios of training and validation sets, as well as different subsets of SNP markers. Genomic Best Linear Unbiased Prediction (GBLUP) and Bayesian Ridge Regression (BRR) combined with expectation maximization (EM) imputation algorithm showed higher PAs and prediction accuracies than Bayesian LASSO (BL). A subset of approximately 4000 markers was sufficient to provide the same PAs and accuracies as the full set of 8719 markers. Furthermore, PAs were similar for both pedigree- and genomic-based estimations, whereas accuracies and heritabilities were slightly higher for pedigree-based estimations. However, prediction accuracies of genomic models were sufficient to achieve a higher selection efficiency per year, varying between 50-87% compared to the traditional pedigree-based selection.

## INTRODUCTION

Scots pine (*Pinus sylvestris* L.) is the most widely distributed pine in the world (Houston Durrant *et al.* 2016; Mátyás *et al.* 2004). It is a highly important commercial species in Europe, particularly in Northern countries (Krakau *et al.* 2013), being the second foremost species for wood production in Sweden (The Swedish National Forest Inventory, 2015). The actual Scots pine breeding program consists of a combination of several selection strategies, all of them based on conventional progeny testing and breeding value prediction based on reliable phenotypic assessments, at age of 10-15 years, and pedigree information, thus a breeding cycle usually takes roughly 21 to 36 years, depending on the testing strategy and mating success (Rosvall *et al.* 2011).

Genomic selection (GS) could potentially reduce the breeding cycle, by shortening field test time through early selections based on GS predictions, and increasing selection intensities with greater genetic gains per unit of time (Crossa *et al.* 2017; Isik 2014; Grattapaglia *et al.* 2018). GS was firstly introduced by Meuwissen *et al.* (2001) and it consisted of using genome-wide marker information to calculate genomic estimated breeding values (GEBV). The major difference between GS and marker assisted selection (MAS) is that there is no need to detect quantitative trait loci (QTL) prior to selection. To perform GS, a training set (TS) of individuals that have been phenotyped and genotyped, generally through single nucleotide polymorphism markers (SNPs), are used to develop prediction models to estimate GEBV, that are validated through a validation set (VS) of individuals, or selection candidates, which are genetically related to the TS and only have marker data for predicting their own breeding values (Grattapaglia and Resende 2011).

Next generation sequencing technologies (NGS) have made it possible to discover thousands of SNPs across the genome and thus make GS a routine application in animal and plant breeding programs (Grattapaglia *et al.* 2018). SNP arrays had been shown as preferable for their reproducibility, manageability and storage logistics, as well as their cost efficiency (Grattapaglia *et al.* 2018). There are still challenges for forest tree species, such as Scots pine, to develop genome-wide SNP panels or exome probe panels because of their large complex genomes, and lack of a reference genome. Therefore, it is attractive to employ alternative genotyping methods such as genotyping-by-sequencing (GBS) (Chen *et al.* 2013; Elshire *et al.* 2011; Dodds *et al.* 2015). GBS uses restriction enzymes to reduce sequencing of complex genomes and uses a barcoding system for multiplex sequencing, which increases its efficiency and reduces the genotyping costs (He *et al.* 2014; Pan *et al.* 2015). GBS can generate very large number of SNPs and produces large amount of missing data. The latter can be solved with the aid of different imputation methods, such as mean imputation (MI), expectation maximization (EM), family-based k-nearest neighbor (kNN-Fam) or singular value decomposition (SVD) (Troyanskaya *et al.* 2001; Dempster *et al.* 1977). EM algorithm was especially designed for GBS data (Endelman 2011; Poland *et al.* 2012). Genomic predictions based on GBS marker information have been successfully studied in animal (Gorjanc *et al.* 2015), crop-(Poland *et al.* 2012; Crossa *et al.* 2013; Jarquin *et al.* 2014) and tree breeding (El-Dien *et al.* 2015; El-Dien *et al.* 2018; Ratcliffe *et al.* 2015).

Accuracy of GS predictions can vary depending on the model selected. Currently different statistical methods are available to estimate GEBV. Genomic best linear unbiased prediction (GBLUP) consists of using the realized relationship matrix (**G** matrix), based on the marker realized kinship relationship, replacing the traditional pedigree numerator relationship matrix (**A** matrix) which is based on coancestry and the infinitesimal model in quantitative genetics, assuming that QTL allelic effects are normally distributed and all have a similar contribution to the genetic variance (Isik *et al.* 2017). On the contrary, most of the Bayesian approaches presume a prior gamma or exponential distribution of QTL allelic effects, thus the variance at each locus can vary (Meuwissen *et al.* 2001). For instance, Bayesian LASSO (BL) assumes that variance follows a Laplace (or double exponential) distribution (Park and Casella 2008). Nevertheless, Bayesian ridge regression (BRR) assigns QTL effects to a multivariate normal prior distribution with a common variance, which is modelled hierarchically through a scaled inverted chi-squared distribution (Perez *et al.* 2010; de los Campos *et al.* 2013; Isik *et al.* 2017).

Although Bayesian approaches may seem more appropriate as they can accommodate different distributions of the allelic effects, the literature on GS in forest trees shows similar results for all models. For instance, Chen *et al.* (2018a) observed similar prediction accuracies when comparing four genomic prediction models (GBLUP, BRR, BL and reproducing kernel hilbert space (RKHS)) in Norway spruce (*Picea abies* (L.). Isik *et al.* (2016) detected similar predictive abilities in maritime pine (*Pinus pinaster* Ait.) comparing GBLUP, BRR and BL prediction models. Although GBLUP and ridge regression BLUP (rrBLUP) were recommended by Tan *et al.* (2017) for their computational advantages, similar predictive abilities were observed for GBLUP, rrBLUP, BL and RKHS, in a *Eucalyptus urophylla* and *E. grandis* hybrid study. In an interior spruce (*Picea engelmannii* × *glauca*) study, Ratcliffe *et al.* (2015) observed similar accuracies for rrBLUP and BayesCπ, which in turn performed better than the generalized ridge regression (GRR), whereas Thistlethwaite *et al.* (2017) observed almost identical predictions with rrBLUP and GRR in Douglas-fir (*Pseudotsuga menziensii* Mirb. (Franco)). On the contrary, Resende *et al.* (2012b) observed better PA for disease resistance in a loblolly pine (*Pinus taeda* L.) study with Bayesian methods when compared with BLUP-based methods. Despite these similar results from different studies carried out so far, it is still important to test the prediction abilities and accuracies of the different genomic prediction models in different species and traits, due to the possible differences in the genetic architecture of the traits.

Among the objective traits of the Scots pine breeding program are: the traditionally existing growth traits, and the recently incorporated wood quality traits (Rosvall and Mullin 2013). The goal of this investigation was to study the prediction power of SNP markers for growth and wood quality traits in Scots pine. The specific objectives were to 1) estimate the predictive ability and prediction accuracy of genomic estimated breeding values (GEBV), 2) compare the efficiency of three different genomic prediction models (GBLUP, BL and BRR) in the estimation of GEBV, 3) study the effect of two different imputation algorithms in the predictive ability and prediction accuracy of GEBV and 4) compare the effect of different numbers of SNPs obtained through GBS in the predictive ability and prediction accuracy of the genomic predictions.

## MATERIALS AND METHODS

### Plant material

In this study a Scots pine full-sib progeny trial (F261, Grundtjärn), belonging to the Swedish tree improvement program at Skogforsk (the Forestry Research Institute of Sweden) was used. The trial consists of 184 full-sib families and 7240 trees (F1-generation), generated from a partial diallel mating design of 40 plus trees (F0-generation) and established in 1971 by Skogforsk as a randomized single tree plot design, divided into 14 post-blocks (Ericsson 1997). A more detailed information on the trial can be found in (Fries 2012). 694 progeny trees (F1) from 183 families were selected for this study, such that the number of trees per family varied from one to seven with an average of four individuals per family.

### Phenotypic data and adjustments

Height (Ht) was measured when the trees were 10 (Ht1) and 30 (Ht2) years old. Diameter at breast height (DBH) was also measured two times, at ages 30 (DBH1) and 36 (DBH2). In 2011, increment cores at breast height were obtained from 694 trees, and processed by Silviscan (Innventia AB, Stockholm, Sweden). From the Silviscan analysis, three traits were used in this study: microfibril angle (MFA), static modulus of elasticity (MOEs) and wood mean density (DEN). In addition, dynamic modulus of elasticity (MOEd) predicted by Hitman ST300 (Fiber-gen, Christchurch, New Zealand) was as well used in the current study. All traits were further described in Hong *et al.* (2014).

The following linear mixed model was applied to reduce the impact of environmental effects for each trait:

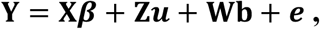

where **Y** is the vector of individual tree observations of a single trait, ***β*** is the vector of fixed effects (intercept), ***u*** is the vector of random effects (post-block and trial design parameters), **b** is a vector of random additive genetic effect of individuals with a normal distribution, 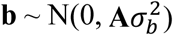, **A** is a matrix of additive genetic effects among individuals, 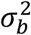 is the additive genetic variance and and ***e*** is the vector of residuals. **X**, **Z** and **W** are the incidence matrices for ***β*, u** and ***b***, respectively.

Adjusted values were obtained for MFA, MOEs, DEN and MOEd, by removing the variation of the experimental design features and post-block effects. For growth traits (Ht1, Ht2 and DBH1 and DBH2), spatial adjustments were performed using the row and column coordinates in the trial. For modeling the residual structure, a model was fitted with only the experimental design elements as factors (Dutkowski *et al.* 2006). If the spatial distribution of residuals were non-random for any trait, a second model was fitted, such that the full residual component was structured as

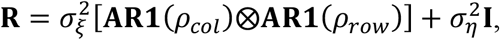

where 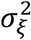 and 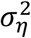 are spatially dependent and independent residual variances, respectively, ⊗ is the Kronecker product of two matrices, and *AR1*(*ρ*_*col*_) and *AR1*(*ρ*_*row*_) represent the first-order autoregressive correlation matrix in the column and row directions, and I denotes the identity matrix (Dutkowski *et al.* 2002; Ivkovic *et al.* 2015; Chen *et al.* 2018b). The adjusted phenotypic data (predicted values of each tree) were used for genomic predictions.

### Genotyping

#### DNA extraction

The commercial NucleoSpin^®^ Plant II kit (Machery-Nagel, Düren, Germany) was used to extract genomic DNA from vegetative buds or needles from the 694 progeny trees and 46 parents. DNA concentration was determined with Qubit^®^ 2.0 fluorometer (Invitrogen, Carlsbad, CA, USA).

#### Genotyping-By-Sequencing (GBS) library preparation

Using 827 samples (replicates included) and *Pst*I high fidelity restriction enzyme (New England Biolabs, MA, USA), three genomic libraries for GBS were prepared following the procedure described in Pan *et al.* (2015). The libraries were sequenced on an Illumina HiSeq 2000 platform at SciLifeLab, Sweden.

#### SNP calling and filtering

Paired-end raw reads of each GBS library were cleaned and demultiplexed by the *process_radtags* module of Stacks v.1.40 (Catchen *et al.* 2011) on the basis of 300 barcodes with 4–8 bp. Cleaned reads of each sample were aligned to the *Pinus taeda* v1.0 (Wegrzyn *et al.* 2014) reference genome, using BWA mem v0.7.15 (Li and Durbin 2010) with default parameters. Alignments were coordinate-sorted and indexed using Samtools v1.5 (Li *et al.* 2009). SNP markers were called using the *mpileup* command of Samtools over all the samples simultaneously, with default parameters, and converted into VCF matrix using BCFtools v0.1.19 (Narasimhan *et al.* 2016). Furthermore, these variants were sorted to keep only high-quality SNPs. Using *vcfutils* in BCFtools with default parameters, the SNPs within 3bp around an indel or with mapping quality < 20 were filtered out; using Vcftools v.0.1.12b (Danecek *et al.* 2011), only SNPs with coverage ≥ 5×, genotype quality (GQ) ≥ 30, genotype calling rate > 20% were kept; using the custom Perl program (ReplicateErrfilter.pl), discordant genotypes of 66 replicated samples were detected and the SNP sites with ≥ 3 replicate errors were filtered out. After this step, 24,152 informative SNP markers were retained.

#### Imputation of missing genotypic data

Missing genotypic data were imputed with TASSEL 5 (Bradbury *et al.* 2007) using LD K-nearest neighbor (Money *et al.* 2015) as a baseline method. After this imputation, two extra imputations were performed to compare their prediction accuracies; random (RND) imputation with the *codeGeno* function in synbreed package (Wimmer *et al.* 2012) in R (R Core Team 2016) and imputation with the expectation maximization (EM) algorithm by the *A.mat* function implemented in rrBLUP package (Endelman 2011) in R. A total of 15,537 and 15,433 SNPs with minor allele frequency (MAF) lower than 1% and with a missing data threshold lower than 10% were removed using RND and EM imputation methods, respectively.

### Statistical analysis for genomic predictions

Among all the available approaches to perform genomic predictions we selected genomic best linear unbiased prediction (GBLUP), Bayesian ridge regression (BRR) and Bayesian LASSO (BL) regression, to estimate genomic breeding values (GEBV) and to evaluate the ability to predict them. ABLUP and GBLUP calculations were performed in ASReml 4.1. (Gilmour *et al.* 2015) whereas BRR and BL were implemented using the *BGLR* function from the BGLR package in R (Perez and de los Campos 2014).

#### ABLUP and GBLUP

Estimated breeding values (EBV) and GEBV were predicted using the following model

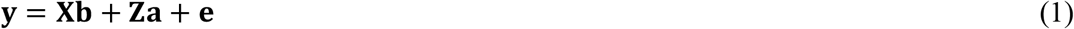

where **y** is the vector of the adjusted phenotypic data for each trait, **b** is the vector of fixed effects (mean), **a** is the vector of random effects and **e** is the vector of residual effects, which is assumed to follow a normal distribution as 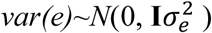, where 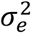 is the residual variance and **I** is the identity matrix. **X** and **Z** are the incident matrices of **b** and **a**.

ABLUP is the method traditionally used to predict EBV, based on pedigree relationships between individuals. In ABLUP, the vector **a** (additive genetic effects) from Eq.1 is assumed to follow a normal distribution with expectations of 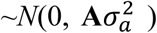, where 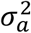 is the additive genetic variance and **A** is the additive numerator relationship matrix.

GBLUP was performed using Eq.1. This method is derived from ABLUP but differs in that the **A** matrix in now substituted by a genomic relationship matrix, known as realized relationship matrix (**G** matrix), estimated according to VanRaden (2008).

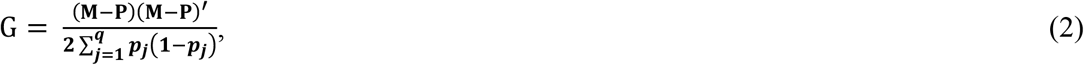

where **M** is the matrix of genotyped samples, **P** is the matrix of allele frequencies with the jth column given by 2(*p*_*j*_ – 0.5), where *p*_*j*_ is the observed allele frequencies of the genotyped samples. The elements of **M** were coded as 0, 1 and 2 (i.e., the number of minor alleles) for the estimation of the **G** matrix with function *kin* from the synbreed package in R in the case of RND imputed data and with the function A.mat from the rrBLUP package in R, for the EM imputed data. The **a** effects from Eq.1, were assumed to follow a normal distribution with expectations of 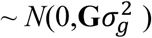, where 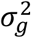 is the genetic variance explained by the markers effect and **G** is the realized relationship matrix, in GBLUP (Isik *et al.* 2017).

#### Bayesian Ridge Regression (BRR)

In BRR, vector **a** from Eq.1 is assigned a multivariate normal prior distribution with a common variance to all marker effects, that is 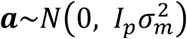, where *p* is the number of markers, 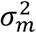 is the unknown genetic variance which is contributed by each marker and assigned as 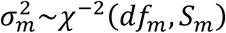, where *df*_*m*_ is degrees of freedom and *S*_*m*_ is the scale parameter. Residual variance is assigned as 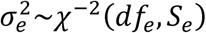, with *df*_*e*_ degrees of freedom and scale parameter for residual variance *S*_*e*_ (Perez et al. 2010).

#### Bayesian LASSO (BL) regression

BL method assumes that vector **a** from Eq.1 follows a hierarchical prior distribution with 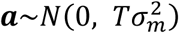, where 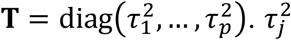 is assigned as 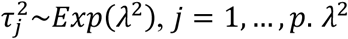 is assigned as *λ*^2^~*Gamma*(*r*, *δ*). Finally, the residual variance is assigned as 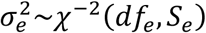, where *df*_*e*_ is degrees of freedom and *S*_*e*_ is the scale parameter for residual variance (Park and Casella 2008).

#### Model convergence and prior sensitivity analysis

Bayesian algorithms were extended using Gibbs sampling for estimation of variance components. The Gibbs sampler was run for 20,000 iterations with a burn-in of 1,000 iterations and a thinning interval of 100. The convergence of the posterior distribution was verified using trace plots. Flat priors were given to all models.

#### Cross validation, prediction accuracy and predictive ability of the models

We performed 10-fold cross-validation, i.e., 90% of individuals in the training set (TS) and 10% in the validation set (VS), for all traits and models (ABLUP, GBLUP, BRR and BL), except to test the different sizes of TS and VS. In addition, for each of the genomic prediction models, two different imputation methods (EM and RND) were evaluated.

For the Bayesian methods, GEBV in the validation set (VS) were estimated as,

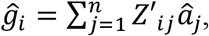

where Z′_ij_ is the indicator covariate (−1, 0, 1) for the *i*^th^ tree at the *j*^th^ locus and 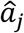 is the estimated effect at the *j*^th^ locus.

Models were evaluated based on their predictive ability (PA) and prediction accuracy (Accuracy). In our study, PA was defined as the Pearson product-moment correlation between the cross-validated GEBVs and the adjusted phenotypes (y) from Eq. 1, i.e., *r*(*GEBV*, **y**) and Accuracy was defined as the Pearson product-moment correlation between the cross-validated GEBVs and the EBVs estimated from ABLUP using all adjusted phenotypes, i.e., *r*(*GEBV*, *EBV*).

#### Effect of the relative size on training and validation sets

The effect on the PA and prediction accuracy, of five different size ratio of TS and VS, was evaluated. The relative size of TS and VS were established dividing the 694 individuals in five different proportions of TS/VS. That is 90%, 80%, 70%, 60% and 50% for TS and the rest as VS. For each trait and each of the 20 models, 10 replicates were performed.

#### Effect of marker number on accuracies

Due to the better predictions obtained with the BRR-EM model from cross-validation results, BRR-EM model was selected to test the effect of the number of SNPs on the PA and prediction accuracy. From all available SNPs, we randomly selected 14 sets of SNPs.

#### Heritability estimation

Pedigree-based narrow sense-heritability 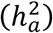 and genomic narrow-sense heritability 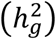 were estimated as

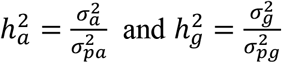

where 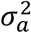 and 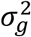 are the pedigree- and genomic-based additive genetic variances and 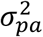 and 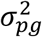 are phenotypic variances estimated using ABLUP and GBLUP, respectively.

#### Relative selection efficiency of GS

Assuming that selection response is inversely proportional to the length of the breeding cycle (Grattapaglia and Resende 2011), the relative efficiency (*RE*) of GS to the traditional pedigree-based selection (TPS) can be estimated as

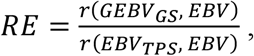

consequently the *RE* per year (*RE/year*) can be estimated as

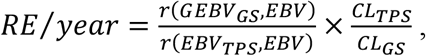

where *CL*_*TPS*_ and *CL*_*GS*_ are the breeding cycle lengths of TPS and GS, respectively.

In order to estimate RE, we assumed that with GS approaches the cycle could be reduced by 50%.

### Data availability

The data sets used in this study are available as File S1 and File S2, in the supplementary material for Calleja-Rodriguez et al. 2019 (link figshare here).

## RESULTS

### Prediction accuracy and predictive ability of the different models

PAs and prediction accuracies from the 10-fold cross-validation were obtained for each model (ABLUP, GBLUP, BRR and BL) and imputation method (Table 1). ABLUP performed best in terms of prediction accuracy. Among the genomic prediction models, different models produced higher accuracies for various traits. There was no single genomic prediction model that fit to all the traits best. In the case of PAs, ABLUP did not showed the highest PA for almost any of the traits. Depending on the trait, the superiority of the models varied for PAs. ABLUP showed higher PA for DEN (0.41); however, it was only slightly higher than PAs obtained with most other models (0.40 in all cases).

**Table 1.** Predictive ability (PA) and prediction accuracy (Accuracy) of each model and trait, ± standard errors.

| Model | Type | Traits |  |  |  |  |  |  |  |
| --- | --- | --- | --- | --- | --- | --- | --- | --- | --- |
|  |  | Ht1 | Ht2 | DBH1 | DBH2 | MFA | MOEs | DEN | MOEd |
| ABLUP | PA | $0.20 \pm 0.01$ | $0.38 \pm 0.000$ | $0.26 \pm 0.003$ | $0.23 \pm 0.01$ | $0.30 \pm 0.001$ | $0.39 \pm 0.01$ | $0.41 \pm 0.01$ | $0.44 \pm 0.02$ |
| | Accuracy | $0.83 \pm 0.01$ | $0.81 \pm 0.000$ | $0.83 \pm 0.003$ | $0.84 \pm 0.01$ | $0.83 \pm 0.001$ | $0.75 \pm 0.01$ | $0.81 \pm 0.01$ | $0.82 \pm 0.01$ |
| GBLUP-EM | PA | $0.20 \pm 0.01$ | $0.39 \pm 0.001$ | $0.26 \pm 0.002$ | $0.26 \pm 0.01$ | $0.29 \pm 0.002$ | $0.39 \pm 0.01$ | $0.40 \pm 0.01$ | $0.41 \pm 0.02$ |
| | Accuracy | $0.69 \pm 0.02$ | $0.75 \pm 0.002$ | $0.73 \pm 0.001$ | $0.74 \pm 0.01$ | $0.73 \pm 0.003$ | $0.69 \pm 0.01$ | $0.73 \pm 0.01$ | $0.74 \pm 0.01$ |
| GBLUP-RND | PA | $0.19 \pm 0.003$ | $0.38 \pm 0.000$ | $0.25 \pm 0.000$ | $0.25 \pm 0.01$ | $0.28 \pm 0.002$ | $0.37 \pm 0.02$ | $0.38 \pm 0.02$ | $0.40 \pm 0.02$ |
| | Accuracy | $0.67 \pm 0.004$ | $0.74 \pm 0.002$ | $0.71 \pm 0.002$ | $0.72 \pm 0.01$ | $0.71 \pm 0.003$ | $0.67 \pm 0.02$ | $0.71 \pm 0.01$ | $0.72 \pm 0.01$ |
| BL-EM | PA | $0.15 \pm 0.04$ | $0.39 \pm 0.02$ | $0.22 \pm 0.02$ | $0.30 \pm 0.04$ | $0.33 \pm 0.03$ | $0.36 \pm 0.03$ | $0.32 \pm 0.02$ | $0.40 \pm 0.03$ |
| | Accuracy | $0.66 \pm 0.03$ | $0.74 \pm 0.01$ | $0.70 \pm 0.02$ | $0.75 \pm 0.02$ | $0.76 \pm 0.02$ | $0.67 \pm 0.02$ | $0.69 \pm 0.01$ | $0.71 \pm 0.02$ |
| BL-RND | PA | $0.26 \pm 0.03$ | $0.36 \pm 0.04$ | $0.26 \pm 0.02$ | $0.26 \pm 0.02$ | $0.28 \pm 0.05$ | $0.34 \pm 0.03$ | $0.40 \pm 0.02$ | $0.41 \pm 0.03$ |
| | Accuracy | $0.69 \pm 0.02$ | $0.73 \pm 0.02$ | $0.71 \pm 0.01$ | $0.72 \pm 0.01$ | $0.68 \pm 0.03$ | $0.65 \pm 0.02$ | $0.71 \pm 0.01$ | $0.72 \pm 0.02$ |
| BRR-EM | PA | $0.18 \pm 0.04$ | $0.41 \pm 0.03$ | $0.25 \pm 0.05$ | $0.27 \pm 0.03$ | $0.33 \pm 0.04$ | $0.42 \pm 0.03$ | $0.40 \pm 0.03$ | $0.46 \pm 0.02$ |
| | Accuracy | $0.65 \pm 0.03$ | $0.77 \pm 0.02$ | $0.72 \pm 0.01$ | $0.75 \pm 0.01$ | $0.73 \pm 0.03$ | $0.70 \pm 0.02$ | $0.72 \pm 0.02$ | $0.76 \pm 0.01$ |
| BRR-RND | PA | $0.24 \pm 0.02$ | $0.39 \pm 0.03$ | $0.21 \pm 0.02$ | $0.24 \pm 0.03$ | $0.27 \pm 0.03$ | $0.40 \pm 0.04$ | $0.40 \pm 0.03$ | $0.45 \pm 0.04$ |
| | Accuracy | $0.72 \pm 0.02$ | $0.75 \pm 0.02$ | $0.70 \pm 0.02$ | $0.74 \pm 0.01$ | $0.73 \pm 0.02$ | $0.68 \pm 0.02$ | $0.72 \pm 0.01$ | $0.75 \pm 0.02$ |

EM and RND denote expectation maximization and random imputation methods, respectively. ABLUP and GBLUP denote pedigree and genomic best linear unbiased predictions, respectively whereas BRR and BL denote Bayesian ridge regression and Bayesian lasso respectively.

In summary, although the best accuracies were observed with ABLUP for all traits, genomic prediction models produced higher PAs for all traits. Moreover, all the genomic prediction models showed similar PAs and prediction accuracies for all traits, being slightly higher when EM imputation method was combined with GBLUP, BRR or BL.

### Relative size effect of the training and validation sets

To test the size effect of different ratios of TS and VS, EM imputation method was used, in combination with ABLUP, GBLUP and BRR since it showed the best PAs and accuracies in our previous 10-fold cross validation.

All three models showed a similar but increasing patterns of PA for different traits with the increase of TS percentages (Fig. 1A). GBLUP and ABLUP showed the highest PAs for almost all traits, when 70% of the individuals were assigned to the TS; however, BRR needed a higher percentage of individuals assigned to the TS to reach the highest PA.

**Fig.1.**
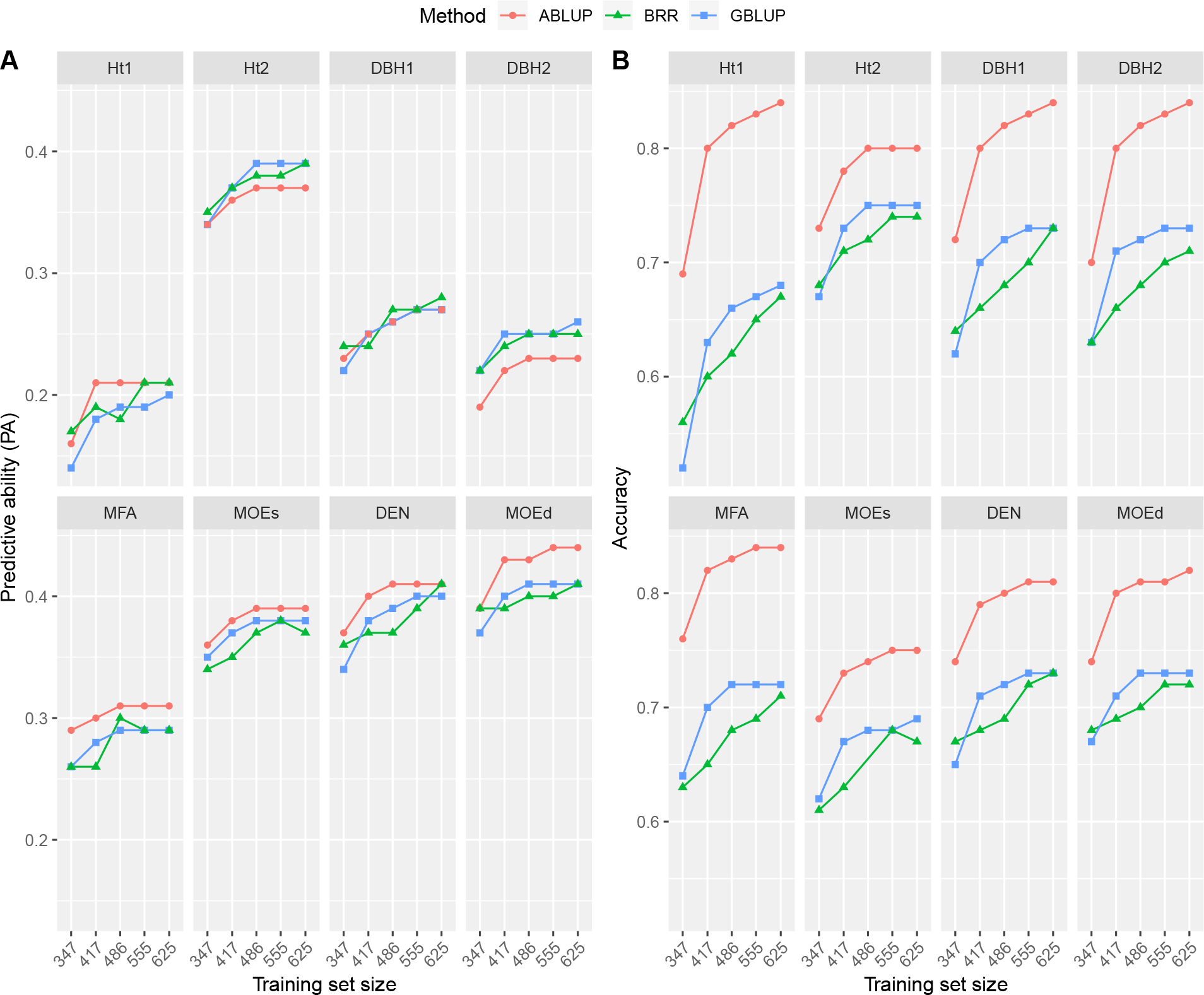
A) Predictive ability (PA) and B) prediction accuracy (Accuracy) of the genomic prediction models for different sizes of training and validation sets.

**Fig. 1.**
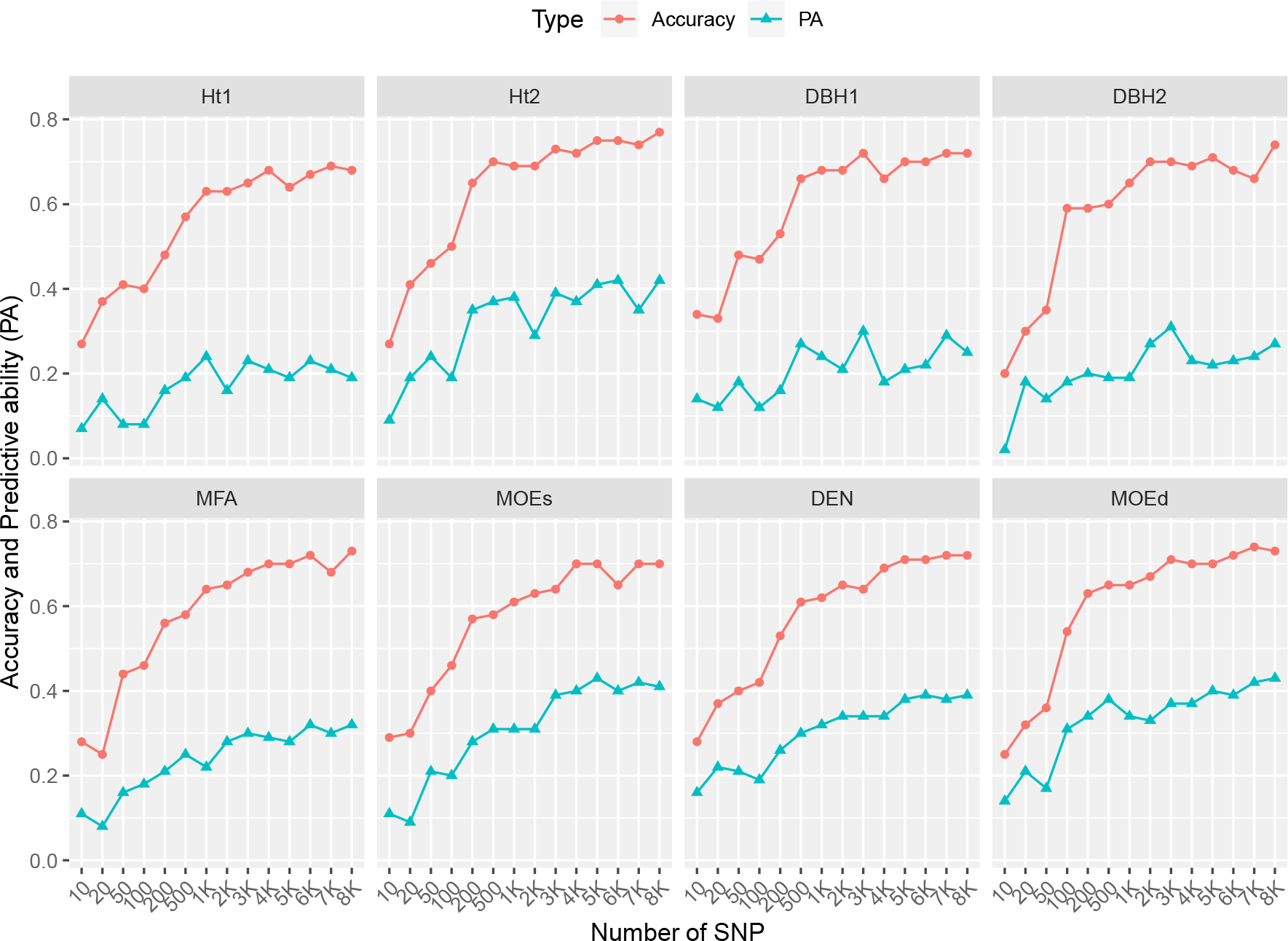
Prediction accuracy (Accuracy) and predictive ability (PA) of Bayesian Ridge Regression prediction model for 14 different subsets of SNPs (10, 20, 50, 100, 200, 500, 1000, 2000, 3000, 4000, 5000, 6000, 7000 and 8719 SNPs).

Among the three methods, ABLUP had the best prediction accuracies for all eight traits under all TS ratios (Fig. 1B). BRR and GBLUP showed similar accuracies. To reach the highest prediction accuracies, 80-90% of individuals in the TS were needed for all traits for BRR method, whereas GBLUP needed a subsample pf 70% or 80% individuals as TS for almost all traits. The computational time needed to perform the analysis as the subset of individuals increased, was substantially longer with Bayesian models.

In brief, the sensitivity analysis suggested that using about 70-80% of individuals sampled from the studied population would produce similar PA and accuracy as the full sample size, for the growth and wood quality traits.

### Effect of increasing number of marker on accuracies

The impact of the different subsets of SNPs was tested based on BRR-EM model that was the model with higher PA and accuracy from the previous 10-fold cross-validation. Accuracies and PAs increased for all traits as the number of SNPs increased (Fig. 2). However, for almost all traits, the greatest increase on prediction accuracy was attained when the subset of markers was 1000 SNPs. Accuracy continued slightly increasing, for all traits with subsets of SNPs higher than 1K, but the increase slowed after 3K – 4K SNPs, reaching the maximum accuracies at 3K for DBH1, 4K for Ht1 and MOEs, 7K for DEN and MOEd, and 8K for Ht2, DBH2 and MFA.

**Fig. 2.**
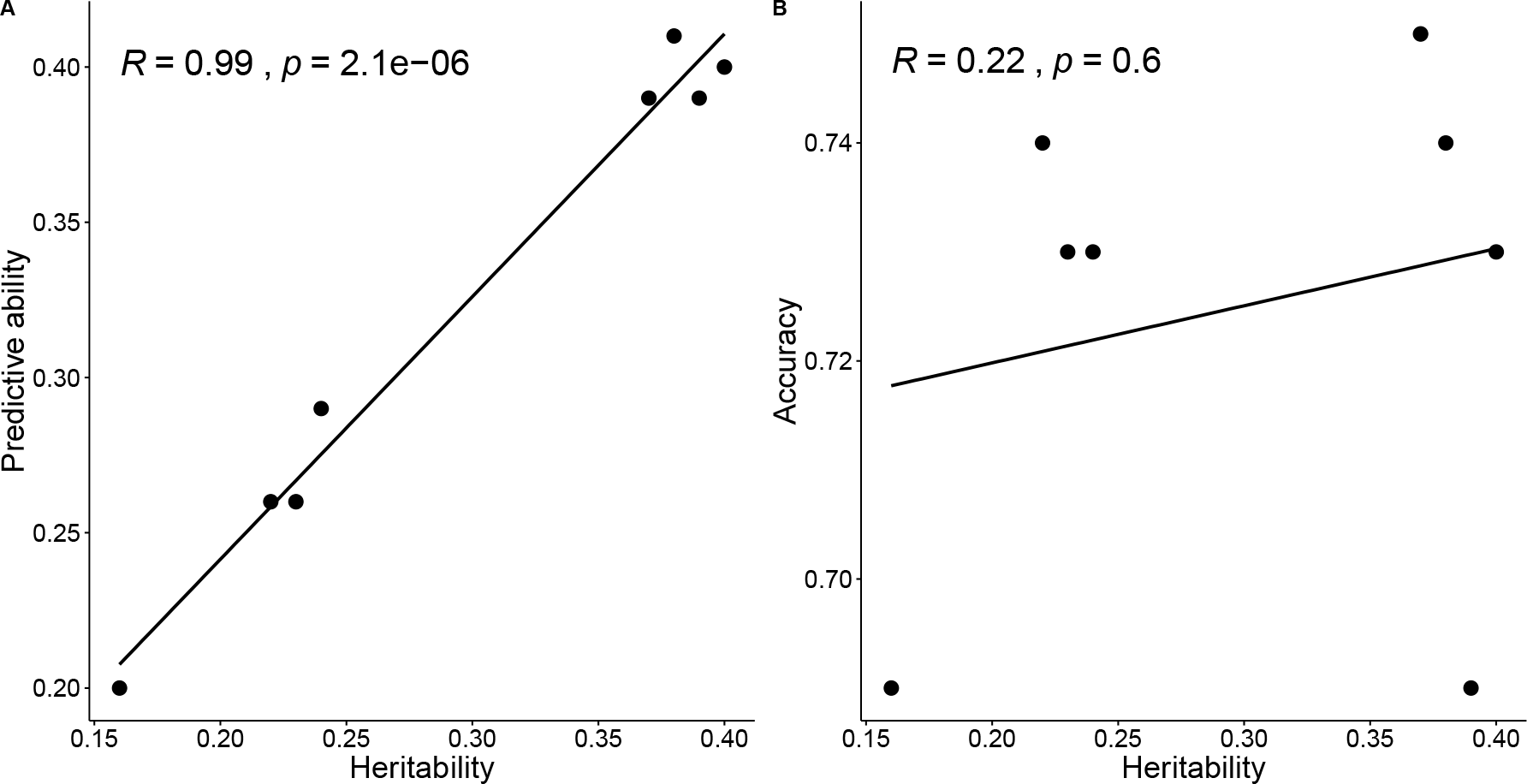
A) Regression between Predictive ability and trait heritabilities. B) Regression between predictive accuracy (Accuracy) and trait heritabilities. Trait heritabilities were estimated with GBLUP-EM model.

PA followed a similar pattern; however, it decreased at a subset of 2K SNPs for Ht1, Ht2 and DBH1 to continue increasing until a subset of 3K SNPs where it stagnated until it reached the maximum of 8719 SNPs. For DBH2, PA decreased at a subset of 4K SNPs and kept constant for the following subset of SNPs. The PA of wood traits showed an increase trend as the number of SNPs rise up, until they reach a plateau at around a subset of SNPs that vary from 4K to 6K depending on the trait. In short, from the subset of 3K-4K SNPs we did not detect any considerable increase in the accuracies and PA of any of the traits except MFA and MOEs for which we detected an increase at the subset of 2K SNPs that kept more or less constant until the final subset of 8719 SNPs.

### Heritabilities

Narrow sense heritabilities estimates based on ABLUP were higher than those based on GBLUP, except for DBH2 which was higher for GBLUP (Table 2). MOEs showed the same heritability both for ABLUP and GBLUP-EM. GBLUP heritability estimates calculated from the realized relationship matrix derived from EM imputation method were higher than those derived from the RND imputation method, for almost all traits, except Ht1 and MOEd. Standard errors were similar for growth traits regardless the BLUP method used but they were always lower when derived from GBLUP method. Based on GBLUP, we observed that traits with heritability estimates equal or lower than 0.25, such as, Ht1, DBH1, DBH2 or MFA, showed estimates of PA below 0.30, while those with heritabilities of approximately 0.40 (Ht2, MOEs, DEN and MOEd) had PA estimations of about 0.40. Moreover, we detected positive linear correlation between PA and trait heritabilities (r=0.99, p<0.0001), but not between accuracies and heritabilities (r=0.22, p=0.6) (Fig. 3).

**Table 2.** Additive genetic variance 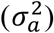 residual variance 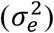 and heritability with standard error (h^2^ ± SE) from ABLUP and GBLUP models.

| Trait | IM | Method | $\sigma_a^2$ | $\sigma_e^2$ | $h^2 \pm \text{SE}$ |
| --- | --- | --- | --- | --- | --- |
| Ht1 | . | ABLUP | 331.3 | 1445.9 | $0.19 \pm 0.06$ |
| | EM | GBLUP | 294.6 | 1504.6 | $0.16 \pm 0.06$ |
| | RND | GBLUP | 305.2 | 1484.3 | $0.17 \pm 0.06$ |
| Ht2 | . | ABLUP | 3827.5 | 5810.3 | $0.40 \pm 0.09$ |
| | EM | GBLUP | 3539.0 | 6170.3 | $0.37 \pm 0.08$ |
| | RND | GBLUP | 3437.0 | 6075.4 | $0.36 \pm 0.08$ |
| DBH1 | . | ABLUP | 147.2 | 460.6 | $0.24 \pm 0.07$ |
| | EM | GBLUP | 144.7 | 473.4 | $0.23 \pm 0.07$ |
| | RND | GBLUP | 133.6 | 475.4 | $0.22 \pm 0.07$ |
| DBH2 | . | ABLUP | 158.8 | 628.7 | $0.20 \pm 0.07$ |
| | EM | GBLUP | 173.4 | 625.6 | $0.22 \pm 0.07$ |
| | RND | GBLUP | 164.4 | 624.2 | $0.21 \pm 0.06$ |
| MFA | . | ABLUP | 4.8 | 12.4 | $0.28 \pm 0.08$ |
| | EM | GBLUP | 4.3 | 13.3 | $0.24 \pm 0.07$ |
| | RND | GBLUP | 4.0 | 13.3 | $0.23 \pm 0.07$ |
| MOEs | . | ABLUP | 1.3 | 2.0 | $0.39 \pm 0.10$ |
| | EM | GBLUP | 1.4 | 2.1 | $0.39 \pm 0.09$ |
| | RND | GBLUP | 1.2 | 2.2 | $0.35 \pm 0.08$ |
| DEN | . | ABLUP | 419.0 | 543.9 | $0.44 \pm 0.10$ |
| | EM | GBLUP | 402.9 | 593.3 | $0.40 \pm 0.08$ |
| | RND | GBLUP | 367.7 | 595.6 | $0.38 \pm 0.08$ |
| MOEd | . | ABLUP | 0.8 | 1.0 | $0.46 \pm 0.10$ |
| | EM | GBLUP | 0.7 | 1.1 | $0.38 \pm 0.08$ |
| | RND | GBLUP | 0.7 | 1.1 | $0.39 \pm 0.08$ |

IM: imputation method. EM and RND denote expectation maximization and random imputations, respectively.

### Relative selection efficiency of GS

The relative genomic selection efficiency (RE) and relative genomic selection efficiency per year (RE/year) were estimated in the genomic selection models, using three models (GBLUP, BRR and BL) and the EM imputation. The Swedish Scots pine breeding cycle combines several selection strategies and we divided in two groups, according to their lengths (Rosvall *et al.* 2011). For the first group of strategies, which is basically seedling backward selection, the cycle length takes up to 36 years. For such strategies, flowering time needs to be included in the cycle length. In order to estimate RE, we assumed that with GS approaches the cycle could be reduced by 50% to 18 years, since 15 years is the starting age for female flowering in Scots pine (Mátyás *et al.* 2004). The cycle length for the second group of strategies (forward selection and open-pollinated backward selection) takes about 21 years and we assumed to shorten this breeding cycle, by 50% as well (11 years) by reducing progeny testing. Both *RE* and *RE*/year for both groups of strategies, were estimated.

The RE/year increased for all traits and models when reducing the breeding cycle by 50% (Table 3). Among the genomic prediction models, highest RE/year were obtained for GBLUP and BRR, which in addition, were slightly higher for the first group of selection strategies than for the second one. The first group of strategies showed RE/year that varied between 66-85% with GBLUP, 57-90% with BRR, and 59-83% with BL, depending on the trait. Within the second group of selection strategies we observed that the RE/year ranged between 59-77% for GBLUP, 50-81% for BRR and 52-75% for BL, again depending on the trait. In summary, for all traits and genomic prediction models, RE/year exceeded 50% when the breeding cycle was reduced by 50%.

**Table 3.** Relative efficiency (RE) and relative efficiency per year (RE/year) of genomic prediction models compared to ABLUP from cross validated models and for each trait.

| Trait | RE |  |  | RE<sup>a</sup>/year |  |  | RE<sup>b</sup>/year |  |  |
| --- | --- | --- | --- | --- | --- | --- | --- | --- | --- |
|  | GBLUP | BRR | BL | GBLUP | BRR | BL | GBLUP | BRR | BL |
| Ht1 | 0.83 | 0.78 | 0.80 | 1.66 | 1.57 | 1.59 | 1.59 | 1.50 | 1.52 |
| Ht2 | 0.93 | 0.95 | 0.91 | 1.85 | 1.90 | 1.83 | 1.77 | 1.81 | 1.74 |
| DBH1 | 0.88 | 0.87 | 0.84 | 1.76 | 1.73 | 1.69 | 1.68 | 1.66 | 1.61 |
| DBH2 | 0.88 | 0.89 | 0.89 | 1.76 | 1.79 | 1.79 | 1.68 | 1.70 | 1.70 |
| MFA | 0.88 | 0.88 | 0.92 | 1.76 | 1.76 | 1.83 | 1.68 | 1.68 | 1.75 |
| MOEs | 0.92 | 0.93 | 0.89 | 1.84 | 1.87 | 1.79 | 1.76 | 1.78 | 1.71 |
| DEN | 0.90 | 0.89 | 0.85 | 1.80 | 1.78 | 1.70 | 1.72 | 1.70 | 1.63 |
| MOEd | 0.90 | 0.93 | 0.87 | 1.80 | 1.85 | 1.73 | 1.72 | 1.77 | 1.65 |

<sup>a</sup> and <sup>b</sup> represent first and second group of selection strategies from the Swedish Scots pine breeding cycle, respectively.
GBLUP, BRR and BL estimates are based on the EM imputation algorithm.

## DISCUSSION

After the genomic selection (GS) concept was proposed in 2001 (Meuwissen et al 2001), genomic prediction studies were initially implemented in dairy cattle. The technology was adopted in crop and tree breeding in the last decade. The execution of GS in animal and crop breeding programs, such as dairy cattle, oat, maize and wheat, increased genetic gains (Meuwissen *et al.* 2016; Crossa *et al.* 2017). Implementation of GS in tree breeding is underway with recent publications in eucalypts (Tan et al 2017), white spruce (Beaulieu et al 2014), black spruce (Lenz et al 2017), interior spruce (Ratcliffe et al. 2015), Norway spruce (Chen et al 2018a), loblolly (Resende et al 2012a, 2012b) and maritime pine (Isik et al 2016). However, genomic prediction studies and new genotyping platforms still need to be developed for many species (Grattapaglia *et al.* 2018). To our knowledge, this is the first genomic prediction study performed in Scots pine.

### Marker imputation for GBS data

For species such as Scots pine, with large and complex genomes (Neale and Kremer 2011) but without a reference genome, and with no SNP chips or exome panels developed, genotyping-by-sequencing (GBS) method is considered as an attractive alternative to perform GS or GWAS studies. When using GBS data, large amounts of missing data are produced, thus filtering and imputation SNPs are critical steps (Dodds *et al.* 2015). In an interior spruce genomic prediction study with GBS data, El-Dien *et al.* (2015) observed that the imputation method used had influence in the quality of predictions and concluded that EM and kNN-Fam imputation methods, provided the highest genomic prediction accuracies. EM was as well the most accurate imputation method in a wheat breeding GS study (Poland *et al.* 2012) with GBS data. Our study support those findings, since among our genomic prediction models we observed more accurate predictions when EM imputation algorithm was used instead of RND imputation, regardless of the genomic prediction model used.

### Accuracy and predictive ability of GS prediction

Traits of interest in tree breeding programs have different genetic architecture; thus, different genomic prediction models to evaluate PA and prediction accuracy must still be studied. Isik *et al.* (2016) observed similar PAs for GBLUP, BRR and BL for growth and stem straightness traits in a two generations genomic prediction study, in maritime pine; however, they found larger bias when BL was used. In a another study with three generations of maritime pine larger bias was detected for ABLUP than for GBLUP or BL (Bartholome *et al.* 2016). Several statistical methods, namely, GBLUP, BRR, BL and reproducing kernel Hilbert space (RKHS), were compared in a Norway spruce study (Chen *et al.* 2018a) where similar prediction accuracies were observed for all of them. rrBLUP, GRR and BayesCπ predictions were compared for interior spruce (Ratcliffe *et al.* 2015), concluding that all methods had similar accuracies although slightly lower for GRR. Congruent with those studies we observed that for wood and growth traits in Scots pine, largest accuracies were obtained with ABLUP for all traits, whereas GBLUP, BL and BRR had similar PAs and accuracies. For instance, accuracies reported in Douglas fir (Thistlethwaite *et al.* 2017), were very similar for height at early age (0.87-0.91) and mature age (0.80 – 0.89), as well as for density (0.94 – 0.96), regardless of the genomic prediction model used, whereas in *Eucalyptus nitents* (Suontama *et al.* 2018), prediction accuracies reported for density (0.74 – 0.79), diameter (0.29 – 0.51) and height (0.29 – 0.51) were slightly lower. Our accuracy estimations for MFA and MOE are similar to those reported for MFA in white spruce (0.71) and MOE in Norway spruce (0.70-0.76), by Beaulieu *et al.* (2014) and Chen *et al.* (2018a), respectively. In addition, PAs for Ht, DBH, MFA and MOE were similar to those reported in Norway spruce, black spruce (*Picea mariana*) or eucalyptus hybrids (Tan *et al.* 2017; Chen *et al.* 2018a; Lenz *et al.* 2017). However, they were slightly lower than those reported for diameter and height in maritime pine (Isik *et al.* 2016).

### Effects of the training and validation set sizes

Our results differed from previous studies which stated that predictive ability and prediction accuracy increased as the size of the training set increased. For instance, Tan *et al.* (2017) detected that PA increased as the TS size increased without reaching any plateau for all models and traits evaluated in *Eucalyptus* hybrids. Similarly, Lenz *et al.* (2017) asserted that accuracy increased as the TS size increased, however after assigning TS of 45% individuals or more, the accuracy increase was not as important. Nevertheless, we found some similarities between other studies, in which the accuracy increased as the TS size increased but reaching a plateau for height when TS reached 80% of individuals and 75% of individuals for wood quality traits (Chen *et al.* 2018a). In the current study, the highest PA and accuracy was obtained when TS size was between 70-80% of the trees, depending on the trait. From those studies, we know, as well, that the number of trees per family have an effect on the GS efficiency; however, we could observe the advantage of applying GS prediction methods, even when the number of trees per family were low.

### Marker number effects

In a general conifer breeding program simulation study, Li and Dungey (2018) detected an increase in the accuracy of GEBV for traits with low and high heritability when the subset of SNPs increased from 7K to 90K, for a training population with 1000 clones from five simulated generations. Moreover, the same pattern was observed in Norway spruce (Chen *et al.* 2018a), where the accuracy increased with number of markers reaching a plateau between 4K and 8K markers. On the contrary, for black spruce, Lenz *et al.* (2017) did not find an remarkable decrease in prediction accuracies when markers were reduced randomly from 5K to 1K; nonetheless, when markers were further reduced to 500, the accuracy decreased dramatically. Tan *et al.* (2017) noted a greater impact of the number of SNPs than their genomic location in the predictive ability, for both GBLUP and RKHS. In the same study, they also observed a stronger reduction in the PA when the subset of SNPs dropped below 5K, and that traits with lower heritabilities were more sensitive to the reduction in the number of SNPs.

The results in this study are in accordance with previous studies (Tan *et al.* 2017; Lorenz *et al.* 2011; Chen *et al.* 2018a) that GBLUP is preferable for large SNP markers datasets, since the Bayesian approaches are computationally demanding, as long as there are no major QTL effects in the study. In the current study 3K to 4K SNP were required to reach a similar efficiency to that achieved when using all 8719 SNPs.

### Heritabilities

Bartholome *et al.* (2016) stated that no clear pattern was detected between accuracy and heritability estimates for maritime pine. Additionally, Grattapaglia and Resende (2011) and Chen *et al.* (2018a) observed that heritability impact on prediction accuracies is relatively insignificant, therefore the former authors recommended that larger training sets should be used for traits with lower heritabilities. Whereas no trend was detected among prediction accuracies and trait heritabilities, we noted a positive linear trend among PA and heritabilities, i.e., traits with lower heritabilities (below 0.25) exhibited the lowest PA while higher PA were detected for traits with moderate heritabilities (above 0.30). This is congruent with the positive correlation between trait heritabilities and PA indicated by Resende *et al.* (2012b) in loblolly pine, that showed a positive trend between trait heritabilities and PA. Similarly, traits with low heritabilities had lower predictive ability in a maritime pine study (Isik *et al.* 2016). Chen *et al.* (2018a) in their Norway spruce study concluded that narrow-sense heritability was more similar to PA than to prediction accuracy, as PA involves both phenotypic and genetic values.

### Relative selection efficiency

A simulation study conducted by Grattapaglia and Resende (2011) showed that when the breeding cycle length was reduced by 50%, the RE/year doubled, and that when the cycle length was reduced by 75% the RE/year reached 3 folds at high marker levels. This theory was confirmed by Resende *et al.* (2012a) that by reducing 50% the loblolly pine breeding cycle, obtained an increase in the RE/year between 53-92% for DBH and 58-112% for Ht, compared to the traditional pedigree-based selection. Similarly RE varied between 106% to 139% for Ht when the breeding cycle length of interior spruce was reduced by 25% (Ratcliffe *et al.* 2015). In Norway spruce, the RE/year of MOE increased between 69 – 83% when the cycle length was also shortened by 50% (Chen *et al.* 2018a). Our results exhibited the same pattern for growth and wood quality traits, with a RE/year ranging between 50 – 90%, with a reduction of the cycle length of 50%.

## CONCLUSIONS

Our results provides an initial perspective of the use of genomic prediction in Scots pine and are encouraging to develop GS strategies for the species. Similar predictive abilities and accuracies among all genomic prediction models were observed, suggesting that the traits are under additive genetic control. Due to both the computational and predictive efficiency, GBLUP was the most effective method to perform genomic predictions for both growth and wood quality traits in Scots pine. The main advantage of GS in Scots pine is the possibility of reducing of the breeding cycle. Our study showed that GS could potentially reduce the breeding cycle by half, and under that assumption, the relative genomic selection efficiency could be as high as 90% depending on the selection strategy and the trait.

The results presented here are based on a relatively small population with a shallow pedigree. More studies using different populations, preferably populations with deeper pedigrees should be carried out to better understand the predictive power of SNP markers for traits with complex inheritance patterns in the species. The predictive power of SNP markers should be tested over two generations as suggested by Isik (2014) because the marker-QTL phase is expected to change once the population undergoes through breeding, due to recombination of homologue chromosomes during the meiosis.

## ACKNOWLEDGEMENTS

We thank to Anders Fries, David Hall, Hong Zhou, Amaryllis Vidalis and René Fernández Uría for field sampling, phenotyping and helping with DNA extractions, Ruiqi Pian for helping with the GBS library preparation. This work was partially funded by Formas (grant number 2014-00171), and also by the Knut and Alice Wallenberg Foundation, and Sweden’s innovation agency Vinnova, through their financial support of the 2^nd^ Research School in Forest Genetics, Biotechnology and Breeding.

